# Regulation of neurofilament length and transport by a dynamic cycle of polymer severing and annealing

**DOI:** 10.1101/2021.01.17.427018

**Authors:** Atsuko Uchida, Juan Peng, Anthony Brown

## Abstract

Neurofilaments are space-filling cytoskeletal polymers that are transported into axons where they accumulate during development to expand axon caliber. We previously described novel severing and end-to-end annealing mechanisms in neurons that alter neurofilament length. To explore the functional significance of neurofilament length, we developed a long-term multi-field time-lapse method to track the movement of fluorescently tagged neurofilaments in axons of cultured neurons for up to 30 minutes. All filaments moved rapidly, but long filaments paused and reversed more often, resulting in little net movement, whereas short filaments moved persistently for long distances, pausing and reversing less often. Long filaments severed more frequently, generating shorter filaments, and short filaments annealed more frequently, generating longer filaments. Thus, neurofilament length is regulated by a dynamic cycle of severing and annealing and this influences neurofilament transport. Site-directed mutagenesis to mimic phosphorylation at four known phosphorylation sites in the head domain of neurofilament protein L generated shorter neurofilaments that moved more frequently. A non-phosphorylatable mutant had the opposite effect. Treatment of cultured neurons with activators of protein kinase A, which phosphorylates three of these sites, increased neurofilament severing. This effect was blocked by the non-phosphorylatable mutant. We propose that focal destabilization of intermediate filaments by N-terminal phosphorylation of their constituent polypeptides at specific locations along their length may be a general enzymatic mechanism for severing this class of cytoskeletal polymers. Our data suggest a novel mechanism for the control of neurofilament transport and accumulation in axons based on regulation of neurofilament polymer length.

**SUMMARY:** Neurofilaments are space-filling cytoskeletal polymers that are transported into axons where they accumulate to expand axon caliber, which is an important determinant of axonal conduction velocity. We reported previously that neurofilaments can lengthen and shorten by novel end-to-end annealing and severing mechanisms. Here, we show that neurofilament annealing and severing are robust phenomena in cultured neurons that act antagonistically to dynamically regulate neurofilament length, which in turn regulates their transport. In addition, we present evidence for a novel enzymatic mechanism of intermediate filament severing based on site-directed phosphorylation of the neurofilament subunit proteins. We propose that modulation of neurofilament length by annealing and severing may be a mechanism for the regulation of neurofilament transport and accumulation in axons.

## INTRODUCTION

Neurofilaments, which are the intermediate filaments of nerve cells, are space-filling polymers that accumulate in axons during development to expand axon caliber and thus increase axonal conduction velocity (Hoffman, 1995; Lariviere and Julien, 2004). These polymers are assembled in the neuronal cell body (Black et al., 1986) and transported into and along axons by the mechanisms of axonal transport (Roy et al., 2000; Wang et al., 2000). Notably, neurofilaments are unique among other known axonally transported cargoes in that they are long flexible protein polymers, measuring just 10 nm in diameter and up to 40 μm or more in length (Fenn et al., 2018a; Fenn et al., 2018b; Yan and Brown, 2005). These polymers move bidirectionally along microtubule tracks powered by kinesin and dynein motors (Francis et al., 2005; He et al., 2005; Uchida et al., 2009). The movement is characterized by short bouts of rapid movement interrupted by prolonged pauses, resulting in an overall slow rate of movement (Brown, 2000). Keratin and vimentin filaments have also been reported to move along microtubule tracks, suggesting that such movements are a general property of intermediate filaments (Leduc and Etienne-Manneville, 2017; Robert et al., 2019).

Studies *in vitro* have shown that intermediate filaments lengthen by joining at their ends, a process known as end-to-end annealing (Herrmann et al., 1999; Mucke et al., 2016; Wickert et al., 2005). Using photobleaching/photoconversion strategies, we demonstrated extensive end-to-end annealing of vimentin filaments and neurofilaments in cell lines (Colakoglu and Brown, 2009) and in primary neurons (Uchida et al., 2013). During those studies, we discovered that neurofilaments can also be severed (Fenn et al., 2018a; Uchida et al., 2013). Subsequently, severing and annealing was reported by others for vimentin filaments (Hookway et al., 2015). End-to-end annealing occurs spontaneously *in vitro* in the absence of accessory factors (Winheim et al., 2011), and likely involves overlap or interdigitation of the staggered coiled coil dimer overhangs at the ends of each intermediate filament (Colakoglu and Brown, 2009; Lopez et al., 2016). However, the mechanism of intermediate filament severing is unknown.

Our discovery that neurofilament length can be altered by severing and annealing has led us to ask whether neurofilament length has any significance for neurofilament function. One possibility is that their length might influence their axonal transport. However, analysis of neurofilament transport on short time scales (seconds or minutes) has revealed no decrease in neurofilament transport velocity with increasing polymer length for filaments ranging from 0.6-42 μm in length (Fenn et al., 2018a). Here, we show that there is in fact a strong dependence of neurofilament transport on filament length, but only on longer time scales (tens of minutes). Long and short filaments both exhibit bouts of rapid movement, but long filaments pause and reverse direction more often. This results in far less net movement of long filaments on long time scales. In addition, we quantify for the first time the rates of annealing and severing in axons and find that these differ with respect to filament length, with long filaments severing more frequently and short filaments annealing more frequently. Our data suggest that neurofilament length and transport are regulated by a dynamic cycle of severing and annealing. We show that this cycle is regulated by phosphorylation and that site-directed phosphorylation may be a mechanism for neurofilament severing.

## METHODS

### Molecular cloning

The rat pNFL-myc, pNFL^S2,55,57,66A^-myc, pNFL^S2,55,57,66D^-myc, pNFL^S2A^, pNFL^S2D^, pNFL^S55A^, pNFL^S55D^, pNFL^S57A^, pNFL^S57D^, pNFL^S66A^ and pNFL^S66D^ cDNA expression constructs were provided by Chris Miller (Yates et al., 2009). The pNFL-myc, pNFL^S2,51,55,66A^-myc and pNFL^S2,55,57,66D^-myc constructs encoded a myc epitope tag linked to the C terminus of rat neurofilament protein L (NFL). To make the pNFL-GFP, pNFL^S2,55,57,66A^-GFP and pNFL^S2,55,57,66D^-GFP fluorescent fusion constructs, we used polymerase chain reaction (PCR) to amplify the NFL sequences in the pNFL-myc, pNFL^S2,51,55,66A^-myc and pNFL^S2,55,57,66D^-myc constructs and then cloned these sequences into the pEGFP-N3 vector (Clontech/Takara Bio) between the EcoR1 and BamH1 restriction sites of the multiple cloning site. The forward and reverse PCR primers were 5’-CGA CTC ACT ATA GGC TAG CCT CGA GAA TTC CTG AGG-3’ and 5’-CGC GGA TCC ATC TTT CTT AGC TTG C-3’ respectively. The resulting vectors coded for GFP linked to the C-terminus of rat NFL by a 5 amino acid linker with the sequence-Gly-Ser-Ile-Ala-Thr-. To make the mouse pNFL-GFP fluorescent fusion construct, we first amplified the mouse NFL coding region by PCR using the mouse pNFL-paGFP construct (Uchida et al., 2013) as a template. The forward and reverse PCR primers were 5’-CAT TGA CGC AAA TGG GCG GTA G-3’ and 5’-CGG GAT CCA TCT TTC TTA GCC AC-3’ respectively. This PCR product was then cloned into the rat pNFL-GFP construct between the NheI and BamH1 sites to replace the rat sequence with the mouse sequence. To make the mouse pNFL^S2,55,57,66A^-GFP and pNFL^S2,55,57,66D^-GFP fluorescent fusion constructs, we used custom gene synthesis (GenScript, NJ, USA) to create 361 bp DNA fragments spanning the first 99 codons of the mouse NFL cDNA (with either the S2,55,57,66A or S2,55,57,66D mutations) flanked by XhoI and PvuI restriction sites. We then cloned these fragments between the XhoI and PvuI restriction sites in the mouse pNFL-GFP construct. The resulting vectors coded for GFP linked to the C-terminus of mouse NFL by a 5 amino acid linker with the sequence-Gly-Ser-Ile-Ala-Thr-, identical to the linker in the rat constructs. All constructs were confirmed by sequencing the entire open reading frame.

### A note about the residue numbering in NFL

There is some inconsistency between the literature and genomic sequence databases concerning the numbering of the amino acid residues in the NFL head domain. NFL was first sequenced by Edman degradation and found to lack the N-terminal methionine and thus start with a serine (Geisler et al., 1985). This was subsequently confirmed by others using mass spectrometry, and the N-terminal serine was found to be acetylated (Betts et al., 1997; Cleverley et al., 1998; Trimpin et al., 2004), which suggests that the N-terminal methionine is removed post-translationally. Thus, the early literature on NFL phosphorylation numbered the phosphorylation sites in the protein starting from the second amino acid residue in the NFL open reading frame and this has continued to the present day (e.g.Giasson et al., 1996; Hashimoto et al., 2000; Nakamura et al., 2000; Yates et al., 2009). To avoid confusion and maintain consistency with the published literature, we chose to retain this numbering. Thus, the serines mutated in the present study are referred to as serines 2, 55, 57 and 66 even though they are actually in positions 3, 56, 58 and 67 of the predicted translation product.

### NFL knockout mice

NFL^+/-^ heterozygous mice (Zhu et al., 1997) were obtained from Don Cleveland at UCSD and maintained heterozygous in our facility by crossing with wild type C57BL/6 mice. NFL^-/-^ homozygotes were obtained in the expected Mendelian proportions by crossing heterozygotes. Genotyping was performed by PCR using tail DNA with forward primer 5’-GCT ATT CGG CTA TGA CTG GGC ACA A-3’ and reverse primer 5’-CGA TGT TTC GCT TGG TCG AAT G −3’, which resulted in amplification of the Neo cassette inserted in Exon 1. The wild type allele was detected using forward primer 5’-GCA ACG ACC TCA AGT CTA TCC GCA-3’ and reverse primer 5’-CTT CGG CGT TCT GCA TGT TCT TGG −3’. The sizes of the amplified PCR products were 366 and 597 bp, respectively.

### Cell culture and transfection

Cortical neurons from mice and rats were cultured using the glial sandwich technique of Kaech and Banker (2006) as modified by Uchida et al. (2016). We used rat glia for both rat and mouse neuronal cultures. To prepare glial cultures, the cerebral cortices of 4 to 6 P0 rats were dissociated and chopped in Ca^++^/Mg^++^-free Hank’s Balanced Salt Solution (HBSS; Invitrogen, Carlsbad, CA). The tissue pieces were dissociated in phosphate-buffered saline (PBS) containing 0.25% [w/v] trypsin (Worthington Biochemical Corp., Lakewood, NJ), 0.1% [w/v] DNase-I (Sigma, St. Louis, MO) and 0.27 mM EDTA (Sigma). The resulting cell suspension was plated in a 75 cm^3^ flask and maintained at 37°C in glial medium consisting of Minimum Essential Medium (Invitrogen) supplemented with 10% [v/v] horse serum (Invitrogen), 0.7% [w/v] glucose (Sigma) and 5 μg/ml gentamicin (Invitrogen), in an atmosphere of 5% CO_2_. After 2 hr, the flask was swirled to remove loosely attached cells and then fed with fresh glial medium. The cells were typically passaged just once about two days before the intended day of use and plated on glass-bottomed dishes coated with poly-D-lysine (Sigma). The glial medium was replaced with neuronal plating medium (see below) on the day of use.

To prepare neuronal cultures, the cerebral cortex of one P0 rat or mouse was dissociated in HBSS and dissociated in PBS containing 0.25% [w/v] Trypsin, 0.27mM EDTA (Sigma) and 0.1% [w/v] DNase-I for each transfection. The resulting cortical neurons were transfected prior to plating by electroporation using an Amaxa Nucleofector™ (Lonza Inc., Walkersville, MD) with the rat neuron Nucleofection kit (VPG-1003) and program G013, or mouse neuron kit (VPG-1001) and program O005. The volume of the cell suspension was 100 μl and the cell density ranged from 4-6 × 10^6^ cells/ml. The DNA consisted of a mixture of 5 μg pNFL-GFP construct, 5 μg pNFM construct and 5 μg EB3-mCherry construct. We used rat constructs when transfecting rat neurons and mouse constructs when transfecting mouse neurons. The cells were then plated onto glass-bottomed dishes coated with poly-D-lysine. Glass coverslips bearing glia (>80% confluency) were suspended cell-side down over the neurons using dots of paraffin wax as spacers. The resulting sandwich cultures were maintained initially at 37°C/5%CO_2_ in neuronal plating medium, which consisted of NbActiv4^™^ (BrainBits; Brewer et al., 2008) supplemented with 5% [v/v] fetal bovine serum (FBS; Thermo Scientific) and 5 μg/ml gentamicin. On the day after plating, this medium was replaced with neuronal culturing medium. The neuronal culturing medium was identical to the plating medium except that it lacked serum and contained 5 μM cytosine arabinoside (AraC; Sigma). Every two days, half the medium was removed and replaced with fresh medium.

Human adrenal adenocarcinoma SW13vim-cells, which lack endogenous vimentin (Sarria et al., 1990), were cultured in DMEM/F12 medium (Thermo Fisher Scientific) supplemented with 5% fetal bovine serum (HyClone, GE Healthcare, Chicago, IL) and 10 μg/ml gentamicin (MilliporeSigma, Burlington, MA). The cells were transfected with Lipofectamine 2000 (Thermo Fisher Scientific) as described previously (Colakoglu and Brown, 2009; Stone et al., 2019).

### Protein kinase A activation

8-bromo cAMP and/or okadaic acid were added directly to the culture medium at final concentrations of 2 mM and 5 nM, respectively. The incubation time prior to imaging was 5 hr for 8-bromo-cAMP and 2 hr okadaic acid. For combined treatment with both reagents, we first added 8-bromo cAMP, incubated for 3 hr, and then added okadaic acid. After incubation for an additional 2 hr, the cells were kept in the medium containing the two reagents and imaged for up to 5 hr.

### Microscopy and imaging

For imaging, the neuronal cell culture medium was replaced with Hibernate-A medium (BrainBits, low fluorescence formulation) supplemented with 2% (v/v) B27 supplement mixture, 0.5 mM Glutamax (Invitrogen) and 37.5 mM NaCl to increase the osmolarity to ~310-320 mOs. To maintain the temperature at 37°C, we used a stage-top incubator with a humidity module (Okolab, Ottaviano, NA, Italy) and an objective heater (Bioptechs, Butler, PA). The cells were observed with a Nikon TiE inverted wide-field epifluorescence microscope equipped with a 100x/1.4NA Plan-Apochromat VC oil immersion objective, SOLA LED light source (Lumencor, Beaverton, OR), and Nikon TI-S-ER motorized stage with X-Y linear encoders. The exciting light was attenuated to 10-15% transmission using the LED light source controller. Images were acquired with no pixel binning using an Andor iXon Ultra 897 EMCCD camera (Andor Technology, Belfast, UK), which has a 512 x 512 array of 16 x 16 μm pixels, and MetaMorph^™^ software (Molecular Devices, Sunnyvale, CA). The GFP and mCherry fluorescence was imaged using an ET-GFP filter cube (model 49002, Chroma Technology, Brattleboro, VT) or an ET-mCherry/Texas Red filter cube (model 49008; Chroma). The movement of GFP-tagged neurofilaments was recorded by acquiring time-lapse movies for a duration of 15 or 30 minutes using 100 ms exposures and 3 s time intervals. For our analyses of the frequency of neurofilament movement, the movies were acquired at a fixed location, which we refer to as fixed-field time-lapse movies. For our analyses of the long-term kinetic behavior of single neurofilaments, we acquired multi-field time-lapse movies by moving the field of view during the acquisition (described below). The movement of EB3-mCherry comets was recorded by acquiring fixed-field time-lapse movies for a duration of 5 minutes using 1 s exposures and 3-5 s time intervals. To compare axon width and neurofilament content in the cortical neuron cultures, overlapping images were acquired along the entire length of multiple axons. Montages were created in the FIJI distribution package of ImageJ (Schindelin et al., 2012) using the Pairwise Stitching plug-in (Preibisch et al., 2009). The images were all acquired with the same exposure and subjected to the same intensity scaling. The axon width was taken to be the width at half height on a linear intensity profile (line width=11 pixels) drawn perpendicular to the long axis of the axon, measured at the mid-point of the axon within each image of the montage, i.e. at approximately 200-300 μm intervals along the axons.

### Multi-field tracking

This method takes advantage of the motorized stage on our microscope and the “Move stage to image position” function in the MetaMorph^™^ software to re-center the filament whenever it approaches the edge of the camera field of view (Uchida et al., 2016). First, we selected an isolated filament to track by observing the acquired images live during time-lapse image acquisition. When the selected filament approached the edge of the camera field of view, we clicked on it. This caused the stage to move automatically to center the filament within the field of view. By repeating this process, it was possible to track single filaments along axons over distances of 1 mm or more. Our time-lapse interval of 3 s was sufficient time to allow the mouse to be clicked, the stage to move, and the new stage position to register before the next image is acquired, thereby allowing continuous time-lapse acquisition of the moving filament. We aimed to follow each filament for 30 min, but we terminated the acquisition earlier than that if the filament reached the end of the axon, entered either the cell body or growth cone, or was lost due to overlap with other filaments. To confirm axon orientation, we acquired a fixed-field timelapse movie of the EB3-mCherry fluorescence and tracked the direction of movement of EB3 comets, which corresponds to the anterograde direction (Fenn et al., 2018a). Axons with ambiguous directionality were excluded from our analysis.

To track neurofilament movement in the multi-field time-lapse movies, the image planes were stitched together using the x-y coordinates of the stage positions to create a multi-field montage from the movie planes (Uchida et al., 2016). The x-y coordinates of one end of the filament were then recorded in successive movie frames using the “TrackPoints” option in the Motion analysis drop-in of the MetaMorph^™^ software. To be consistent, for each filament we selected the end that was leading when the filament was moving in its net direction of movement and tracked that same end throughout the movie. The coordinates and other parameters such as elapsed time, direction, and velocity were logged to a Microsoft Excel spreadsheet via Dynamic Data Exchange for further analysis.

### Analysis of neurofilament movement, severing and annealing

For the multi-field time-lapse movies, all filaments greater than or equal to 10 pixels (1.6 μm) in length were analyzed if they moved a total distance of at least 50 pixels (8 μm). The net velocity was calculated by dividing the net distance moved by each filament during each movie by the elapsed time. Since neurofilaments can exhibit frequent folding behaviors while they are pausing but are almost always fully extended when they move (Fenn et al., 2018b), we only measured filament length during bouts of movement. To avoid motion blurring, we measured the filaments during brief pauses rather than during bouts of rapid movement. To quantify annealing frequency, we only counted events in which the adjoined filaments moved together as one after the putative annealing event. We measured the filaments before and after the putative annealing event to ensure conservation of length. To quantify severing frequency, we only counted events in which the filaments moved before breaking (in order to be sure that it was indeed a single filament) and in which there was clear separation between the resulting fragments (e.g. due to movement of one or both of the fragments). To quantify the frequency of neurofilament movement in fixed-field time-lapse movies, we located the midpoint of the longest gap in the neurofilament array in the field of view and drew a line perpendicular to the axon at that location. We then counted the number of filaments that moved across each line during that movie and divided that number by the duration of the movie. If there were multiple gaps along a single axon, we selected the longest for our analysis; we did not analyze more than one gap per axon.

### Immunofluorescence microscopy

Neurons were fixed with 4% formaldehyde and then extracted with PBS containing 1% [v/v] Triton X-100 and 0.3 M NaCl. After fixation and extraction, the cells were incubated sequentially with a mouse monoclonal specific for phospho-independent NFM (RMO270; Invitrogen, 1:400) and a rabbit polyclonal antibody specific for the myc tag (71D10: Cell Signaling, 1:100). The secondary antibodies were Alexa 488-goat anti-mouse (Invitrogen, 1:200) and Alexa 568-goat anti-rabbit (Invitrogen, 1:200). Coverslips were mounted using ProLong Gold Antifade reagent (Invitrogen). Images were acquired on an Andor Revolution WD spinning disk confocal, which incorporates a Yokogawa CSU-W1 confocal scanning unit, Nikon TiE inverted microscope, 100x Plan Apo 1.0 NA oil immersion objective and Andor iXon Ultra 897 EMCCD camera. SW13vim-cells were fixed with 4% formaldehyde in PBS and immunostained as described above. Images were acquired on the same microscope but using an Andor Neo sCMOS camera.

### Statistical analysis

The relationship between filament length and various transport kinetic parameters was tested using a linear regression model. The relationship between filament length and the probability of annealing and severing events was tested using a logistic regression model. The effects of NFL phosphorylation or activators of PKA on filament length were tested using a linear regression model. Filament lengths were natural log-transformed to better approximate normality of the residuals. The effect of NFL phosphorylation on transport frequency was tested using a Poisson regression model. A two-sided significance level of α=0.05 was used for all tests. The analyses were performed in SAS version 9.4 (SAS, Cary, NC). Statistical significance is represented in the figures as follows: ns (p>0.05), * (p≤0.05), ** (p≤=0.01), *** (p≤0.001), **** (p≤0.0001).

## RESULTS

To visualize neurofilaments, we co-transfected dissociated cortical neurons with GFP-tagged NFM (GFP-NFM) (Wang and Brown, 2001) and EB3-mCherry (to allow assessment of axon orientation). We then observed the cells after 8-10 days in culture. The GFP fusion protein coassembles fully with endogenous neurofilament proteins and the low neurofilament content of these neurons makes it possible to track single moving filaments (Uchida et al., 2016). Our previous time-lapse imaging of neurofilament transport was limited to short-term observations because the filaments moved rapidly out of the camera field of view. To overcome this problem, we used the motorized stage on our microscope to re-center the moving filaments when they approached the edge of the field of view, allowing us to record their movement across multiple fields of view (see Methods). We then used the stage position of each frame in the resulting movies to register the images, creating a movie montage (Movie S1). Using this multi-field timelapse tracking approach, we were able to follow the movement of single neurofilaments along axons for distances of 0.5 mm or more (Fig. 1A).

**Fig. 1.**
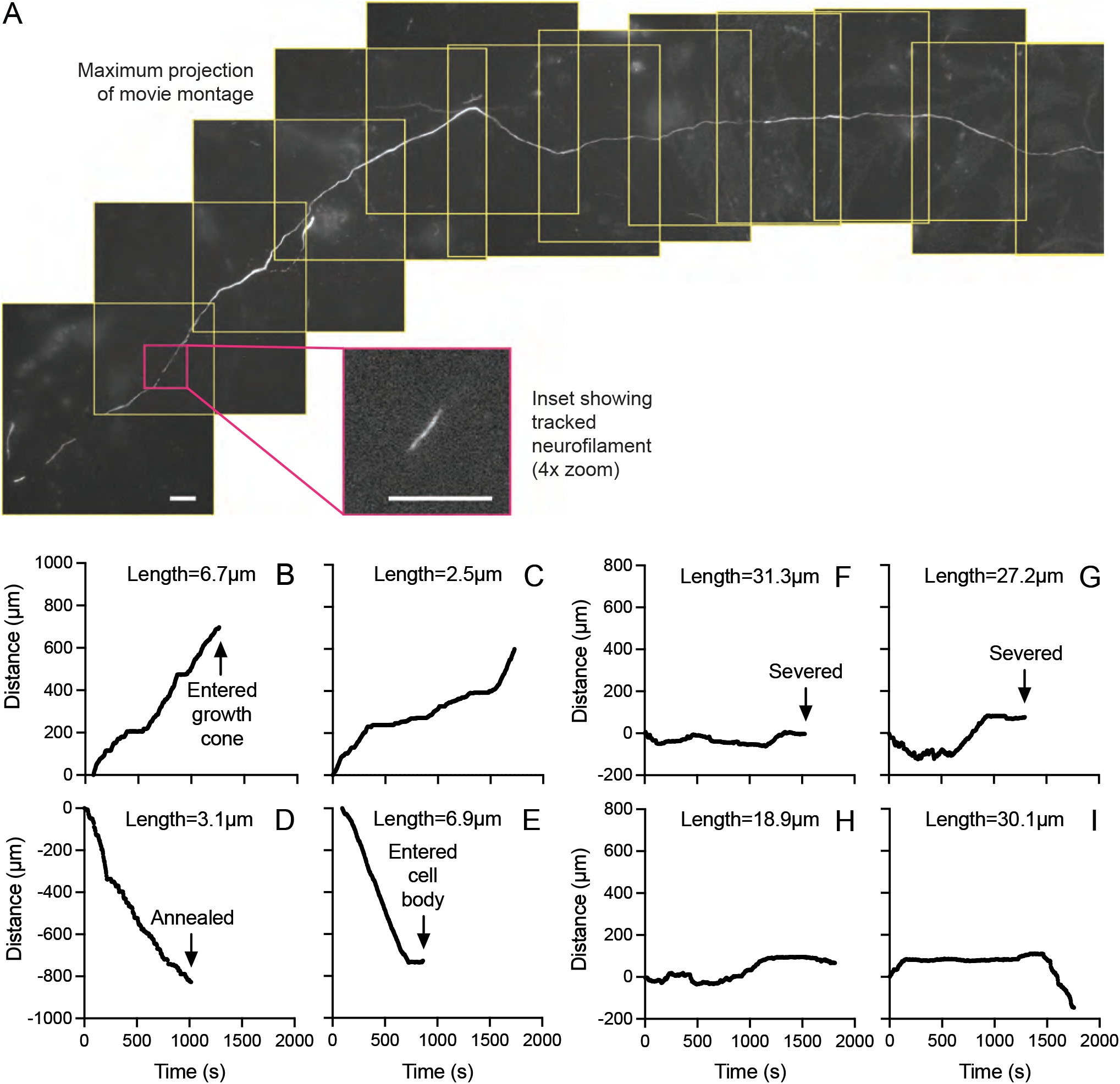
Long-range multi-field tracking of neurofilaments. **A.** Maximum intensity projection of a multi-field time-lapse movie showing the trajectory of a neurofilament along an axon of a neonatal rat cortical neuron after 8 days in culture. The microscope stage was moved 10 times during the movie, allowing the moving filament to be tracked for approximately 420 μm over a period of 12.4 min (248 images acquired at 3 s time intervals). The images were aligned using the encoded microscope stage coordinates to create a movie montage comprised of 11 distinct overlapping fields (yellow boxes). The inset is a region of one image plane at 4x zoom, showing the tracked filament at higher magnification. Scale bar=10 μm in main figure and inset. **B-E.** Representative trajectories of four short filaments (<10 μm in length). **F-I.** Representative trajectories of four long filaments (>15 μm in length). Data from time-lapse movies of neonatal rat cortical neurons after 8-10 days in culture. Distance along the axon is plotted versus time. Anterograde is positive and retrograde is negative. Each filament was tracked for up to 1800 s (30 min). The tracking was terminated early if the filament reached one end of the axon (B,E), annealed with another filament (D), or severed (F,G). Note that the short filaments moved rapidly and persistently in the anterograde or retrograde direction, whereas the long filaments also moved rapidly, but paused and reversed direction more frequently.

### Dependence of transport kinetics on filament length

To investigate the dependence of neurofilament transport on polymer length, we acquired 76 long-term time-lapse movies of moving filaments in neonatal rat cortical neurons after 8-10 days in culture. The resulting movies often also captured the movement of other filaments. Thus, during the subsequent analysis, we tracked all filaments in the movies that met our analysis criteria (see Methods). In total, we obtained trajectories for the movement of 93 filaments, which ranged from 1.3 to 48.3 μm in length. Fifty-five moved in a net anterograde direction and 38 moved in a net retrograde direction.

Fig. 1 B-I shows representative examples of the trajectories of eight filaments, four “short” filaments (2.5-6.9 μm in length) and four “long” filaments (18.9-31.3 μm in length). As we have reported previously, filaments of all lengths moved rapidly on short time scales (seconds to minutes) (Fenn et al., 2018a). However, we observed a marked difference on longer time scales (tens of minutes). Short filaments exhibited a clear directional preference (anterograde or retrograde), and they tended to move persistently in that direction throughout the tracking period, pausing and reversing relatively infrequently. Long filaments were also capable of rapid movement, sometimes for several minutes or more, but they paused more and reversed direction more frequently. As a result, the movement of long filaments often resulted in little net displacement.

To quantify the relationship between transport kinetics and filament length on long time scales, we further analyzed all those filaments that we were able to track for at least 8 min (n=65 filaments). Fig. 2 shows scatter plots of net average transport velocity, percent time pausing, directional persistence and peak velocity versus filament length. Filaments that moved in the retrograde direction had a 0.138 μm/s greater net average velocity compared to those moved in the anterograde direction (p=0.006) (Fig. 2A). This is consistent with previous reports that neurofilament movement is faster in the retrograde direction than in the anterograde direction (Fenn et al., 2018a). However, for both directions regression analysis revealed a marked and statistically significant decrease in net average velocity with increasing filament length (p<0.0001). Quantification of the moving and pausing behavior confirmed that this decrease in net average velocity was due to an increase in the time that the filaments spent pausing (p<0.0001) (Fig. 2B) and a decrease in their directional persistence, i.e. the proportion of the time the filaments spent moving in the direction of their net movement (p<0.0001) (Fig. 2C). There was no statistically significant difference in percent time pausing (p=0.5) or directional persistence (p=0.15) with respect to the direction of movement, but the peak velocity was 1.72 μm/s greater in the retrograde direction than in the anterograde direction (p<0.0001). For every 1 μm increase in filament length, we observed a velocity decrease of 0.016 μm/s, a 1.1% increase in percent time pausing and a 1.5% decrease in directional persistence. In contrast, though there was considerable scatter in the data, the absolute peak velocity exhibited no significant length-dependence in either direction (p=0.32) (Fig. 2D).

**Fig. 2.**
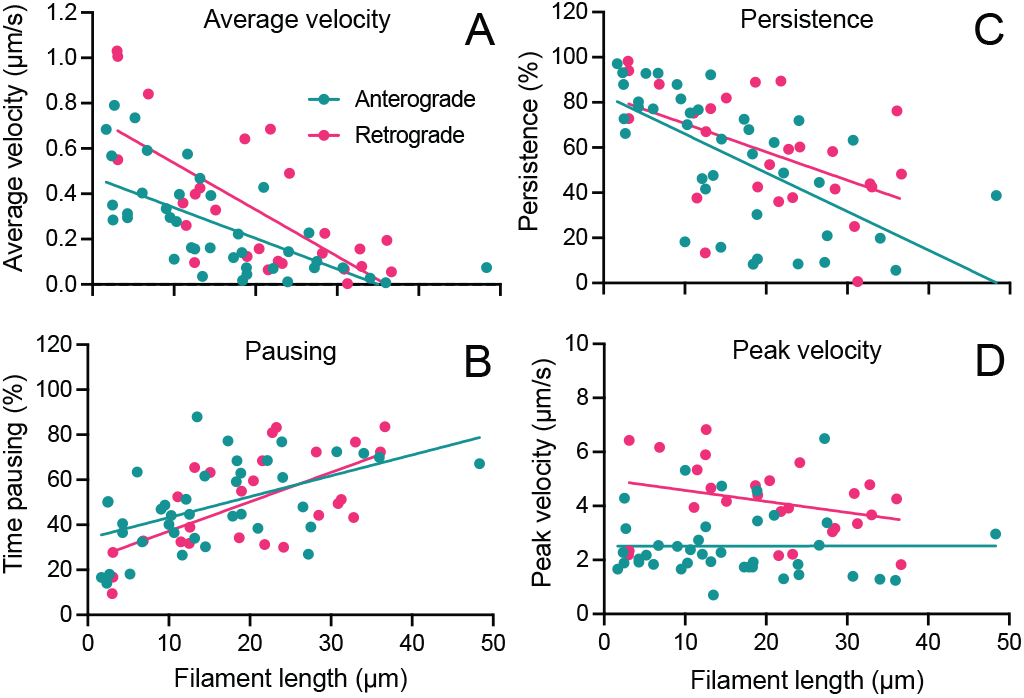
Length-dependence of net neurofilament transport velocity. Scatter plots of the kinetic parameters of neurofilament transport versus filament length for 65 filaments that were tracked for at least 8 min and up to 30 min each in time-lapse movies of neonatal rat cortical neurons after 8-10 days in culture. **A.** Average velocity, which is the net distance moved divided by the duration that the filament was tracked. There was a statistically significant effect of direction (p=0.006) and length (p<0.0001) on velocity. **B.** Percent time pausing. There was a statistically significant effect of length (p<0.0001) but not direction (p=0.497) on pausing. **C.** Directional persistence, which is the net distance moved divided by the total distance moved (irrespective of direction). There was a statistically significant effect of length (p<0.0001) but not direction (p=0.15) on persistence. **D.** Peak velocity, which is the maximum time-lapse interval velocity for each filament. There was a statistically significant effect of direction (p<0.0001) but not length (p=0.32) on peak velocity. The lines show the linear regression for filaments that moved in a net anterograde (teal) or retrograde (magenta) direction. Note that the average net velocity declined with increasing neurofilament length for filaments that moved in either net direction, and this was due to an increase in the percent time pausing and a decrease in the directional persistence of the movement.

### Dependence of annealing and severing frequency on filament length

As mentioned above, the filaments that we tracked sometimes exhibited severing or end-to-end annealing. Examples are shown in Fig. 3 and Movies S2 and S3. To quantify annealing and severing frequency, we counted the total number of annealing and severing events in the 76 movies analyzed above. We also included an additional 21 movies that we excluded from the original data set due to ambiguity in the axon orientation or due to the use of time-lapse intervals of 2 s instead of 3 s. In total, we observed 29 severing events and 15 end-to-end annealing events, corresponding to an average severing rate of 0.9/hr and an average annealing rate of 0.5/hr (141 filaments tracked; 32 hr total tracking time). Thus, approximately one third of the filaments that we tracked exhibited a severing or annealing event. To investigate the effect of filament length, we analyzed the dependence of these rates on the length of the tracked filament (Fig. 3C,D). Analysis using binary logistic regression revealed that there was a statistically significant difference between the average length of filaments that annealed (9.3±7.7 μm, n=15), those that severed (22.1±11.9 μm, n=29) than those that neither annealed or severed (15.6±12.2 μm, n=97) (p=0.0007). For every 1 μm increase in filament length, the probability of severing increased by 4.7% (95% confidence interval: 1.01-1.08) and the probability of annealing decreased by 7.6% (95% confidence interval: 0.86-0.99). Short filaments (<10 μm) exhibited the highest rate of end-to-end annealing (1.24 events/hr) but the lowest rate of severing (0.34 events/hr). In contrast, long filaments (>20 μm) exhibited the lowest rate of annealing (0.15 events/hr) and the highest rate of severing (1.5 events/hr). Thus, the rates of severing and end-to-end annealing exhibited inverse length dependencies.

**Fig. 3.**
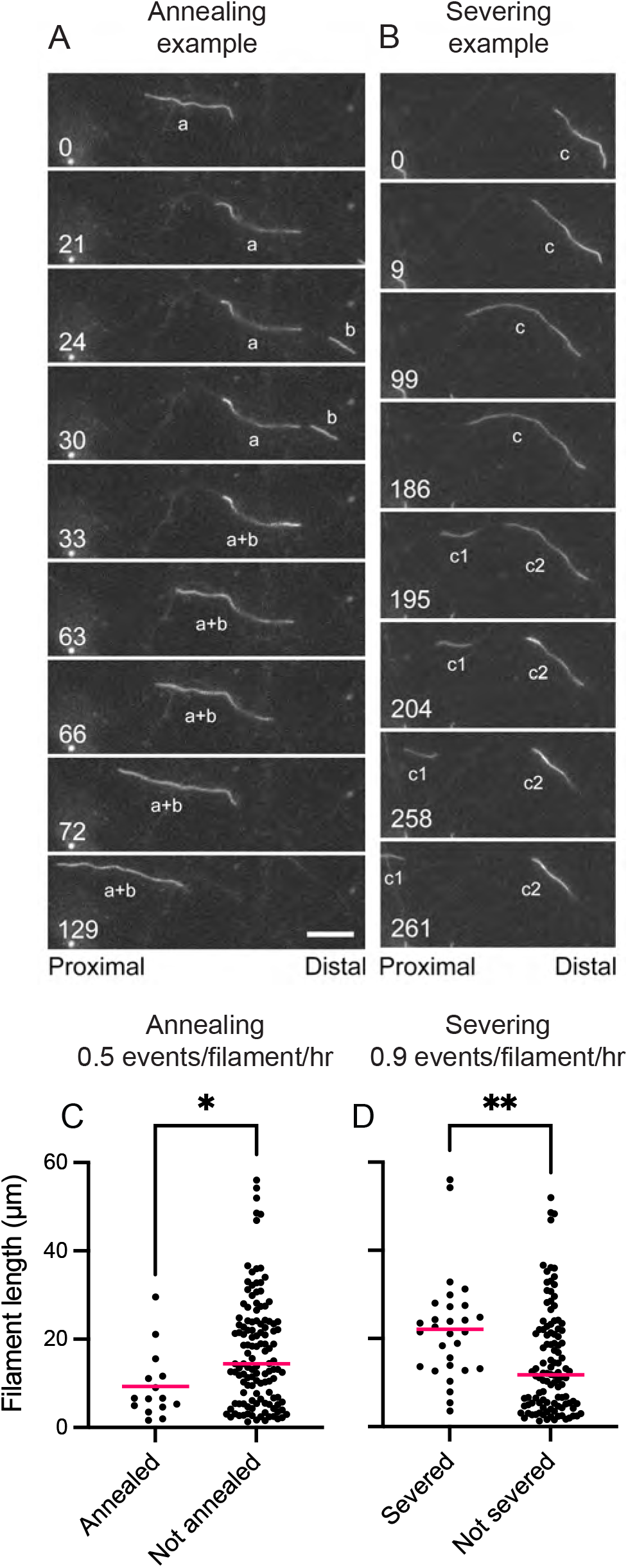
Length-dependence of neurofilament severing and annealing. **A,B.** Excerpts from time-lapse movies of an annealing and a severing event (see Movies S2 and S3, respectively). Time is shown in seconds. In A, filament “a” moved anterogradely initially (t=0-21s) and then paused and folded along part of its length resulting in an increase in brightness at its proximal end. Note that a portion of this filament was out of focus because the axon passed slightly outside the plane of focus at that point. Filament “b” then moved retrogradely into the field of view from the right and overlapped filament “a” (t=33s). The two filaments subsequently annealed and the resulting daughter filament moved retrogradely, stretching out as it did so (t=63-129s). Filament lengths: 20.8 μm (a), 6.3 μm (b), 27.5 μm (a+b). In B, filament “c” was initially paused and partially folded (t=0s) and then stretched out as it moved retrogradely (t=9-99s). The filament then paused (t=99-186s) before severing (t=195s). The leading filament fragment (“c1”) continued to move retrogradely and the trailing fragment (“c2”) folded back on itself (note the increase in brightness in the folded region) and remained paused. Filament lengths: 29 μm (c), 7.1 μm (c1), 21.6 μm (c2). Scale bar=10μm. **C,D.** Dot plots of filament length for filaments that either annealed or did not anneal, and that either severed or did not sever, during the period of observation. Data from 97 movies (total=141 neurofilaments observed for 32 hr; average time-lapse movie duration=13.6 min). In all cases, filament length was taken to be the length before the severing or annealing event. The average annealing and severing frequencies are shown above the graphs. Note that short filaments annealed more frequently than long filaments and that long filaments severed more frequently than short filaments (p=0.024 in C; p=0.0057 in D). The horizontal magenta lines represent the population means.

### Phospho- and dephospho-mimics of NFL have different effects on neurofilament length, transport, severing and annealing

It is known that the intermediate filament protein head domains regulate filament assembly (Gill et al., 1990), and phosphorylation of these domains induces intermediate filament fragmentation and disassembly *in vitro* and *in vivo* in a variety of cell types (Ando et al., 1989; Chou et al., 1996; Eriksson et al., 2004; Hisanaga et al., 1990; Inagaki et al., 1987; Izawa and Inagaki, 2006; Klymkowsky et al., 1991). Thus, phosphorylation of the N-terminal head domains of the neurofilament subunit proteins at loci along neurofilament polymers could be a mechanism for neurofilament severing. To test this hypothesis, we obtained a panel of rat NFL constructs from Chris Miller (King’s College, University of London) in which the four serines in the NFL head domain that have been proposed to be phosphorylated in neurons (serines 2, 55, 57, 62) were mutated to either aspartate or alanine (Yates et al., 2009). Serines 2, 55 and 62 have been reported to be phosphorylated by protein kinase A (Cleverley et al., 1998; Giasson et al., 1996; Nakamura et al., 2000; Sihag and Nixon, 1991; Trimpin et al., 2004), and serine 57 by Rho Kinase and calcium-calmodulin dependent kinase (CaMKII) (Hashimoto et al., 1998; Hashimoto et al., 2000).

Both the NFL^S2,55,57,66A^ (dephospho-mimic) and NFL^S2,55,57,66D^ (phospho-mimic) mutants coassembled with NFM in SW13vim-cells (data not shown). To investigate the effect of NFL phosphorylation on neurofilaments in neurons, we expressed fluorescent fusions of these mutants and of wild type NFL (NFL^S2,55,57,66A^-GFP, NFL^S2,55,57,66D^-GFP or NFL-GFP, respectively) in cultured neonatal rat cortical neurons. In comparison to GFP-NFM, transfection with NFL-GFP alone tended to result in elevated diffuse fluorescence, which is indicative of incomplete incorporation into filaments. We found that this could be avoided by co-transfecting the NFL-GFP fusions with untagged NFM. Under these conditions, both the NFL^S2,55,57,66A^-GFP and NFL^S2,55,57,66D^-GFP constructs incorporated fully into neurofilament polymers throughout the cells (Fig. 4). In contrast to Yates et al. (2009), we did not observe any tendency for the NFL^S2,55,57,66D^ mutant to aggregate.

**Fig. 4.**
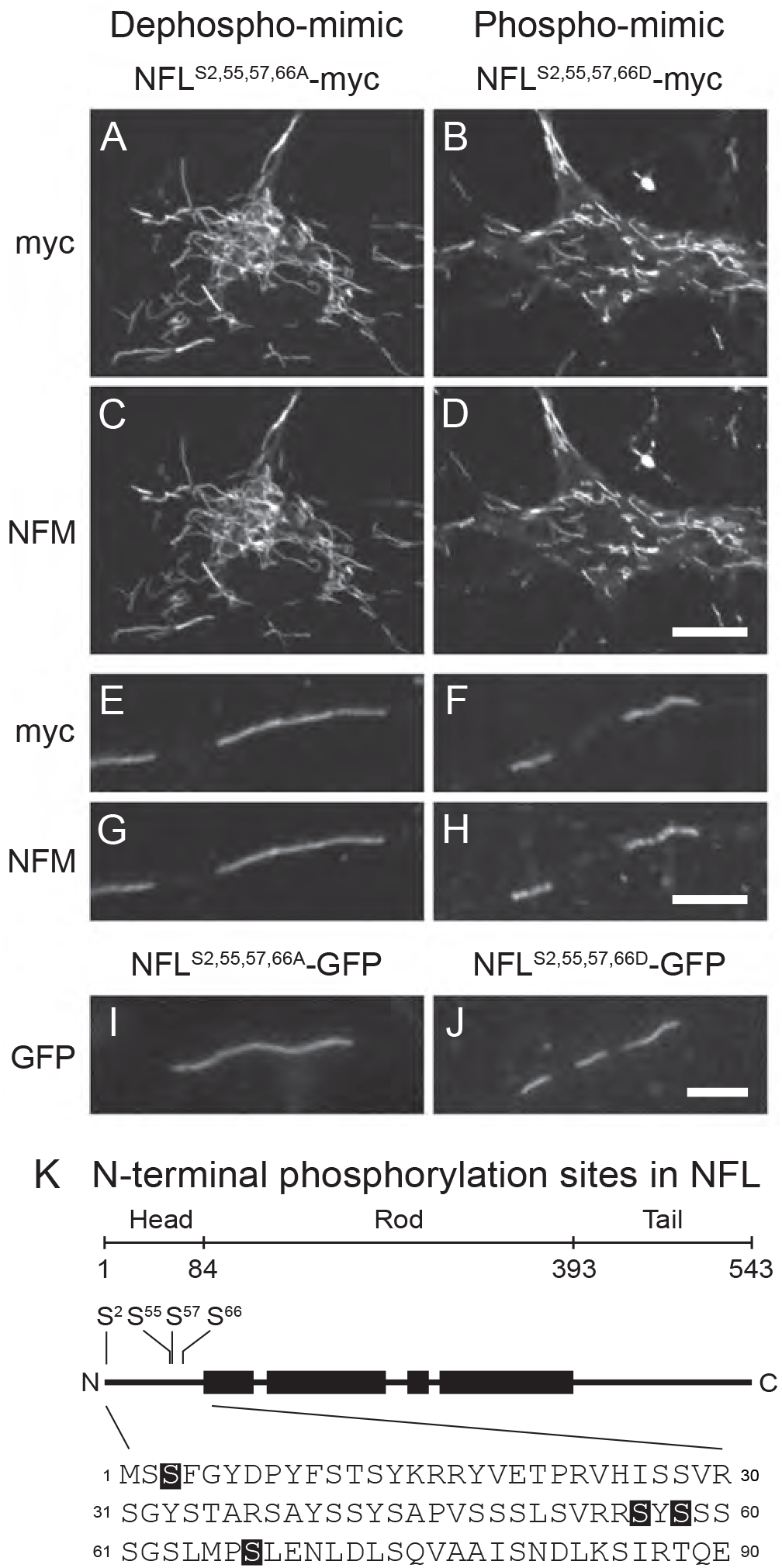
NFL phospho- and dephospho-mimics assemble into neurofilaments. **A-H.** Spinning disk confocal microscopy of neurofilaments in cell bodies (A-D) and axons (E-H) of neonatal rat cortical neurons expressing NFL^S2,55,57,66A^-myc (dephospho-mimic) or NFL^S2,55,57,66D^-myc (phospho-mimic) 10-11 days after plating, immunostained for myc and NFM. The colocalization of myc and NFM confirms that the mutant NFL co-assembled into neurofilaments with NFM. **I,J.** Live wide-field imaging of axonal neurofilaments in axons of neurons expressing NFL^S2,55,57,66A^-GFP or NFL^S2,55,57,66D^-GFP 10 days after plating. Note the uniform incorporation of the mutant proteins along the neurofilaments and the absence of aggregates or diffuse fluorescence, which confirms that the mutant proteins assemble fully into neurofilaments. On average, the neurofilaments containing the phospho-mimic mutant were shorter, which is confirmed by the quantitative analysis in Fig. 6. Scale bars= 5 μm. **K.** Schematic showing the location of the mutated serine residues in the N-terminal head domain of the NFL protein. Note that the numbering of the residues ignores the N-terminal methionine (see explanation in Methods).

First, we investigated the neurofilament length distribution and transport frequency in fixed-field time-lapse movies of neurons expressing wild type NFL-GFP, NFL^S2,55,57,66D^-GFP (phosphomimic) and NFL^S2,55,57,66A^-GFP (dephospho-mimic) (Fig. 5A-D). We observed statistically significant interactions for both length and frequency with respect to age in culture (p<0.0001), which is indicative of different change patterns among the three groups. For neurons expressing wild type NFL, the average filament length increased with age in culture from 6.3 μm at 8 days to 9.2 μm at 11 days, and the average frequency decreased from 7.9 filaments/axon/hr at 8 days to 2.3 filaments/axon/hr at 11 days. Thus, neurofilaments became longer and less mobile as these neurons matured. Filaments containing the phospho-mimic NFL were shorter and increased only slightly in length, from 5.1 μm at 8 days to 5.8 μm at 11 days (p=0.023), but showed a similar frequency of movement to the wild type that decreased from 8.2 filaments/axon/hr at 8 days to 4.6 filaments/axon/hr at 10 days (p<0.0001). In contrast, filaments containing the dephospho-mimic NFL decreased in length from 13.5 μm at 8 days to 9.1 μm at 10 days (p=0.0042) and then increasing slightly to 9.7 μm at 11 days, though this was not statistically significant (p=0.08). The frequency of movement of these filaments was significantly lower than for filaments containing wild type and phospho-mimic NFL (p<0.0001), but also decreased over time from 3.1 filaments/axon/hr at 8 days to 2.1 filaments/axon/hr at 10 days (p=0.016). Interestingly, we also observed that in older cultures (after 16-21 days in vitro), the neurofilament content was higher in axons expressing the dephospho-mimic NFL^S2,55,57,66A^-GFP than in axons expressing the phospho-mimic NFL^S2,55,57,66D^-GFP, resulting in a significant increase in axon width (Fig. 5E,F). Thus, phosphorylation of these sites regulates neurofilament length, which in turn regulates neurofilament transport and accumulation during axonal maturation.

**Fig. 5.**
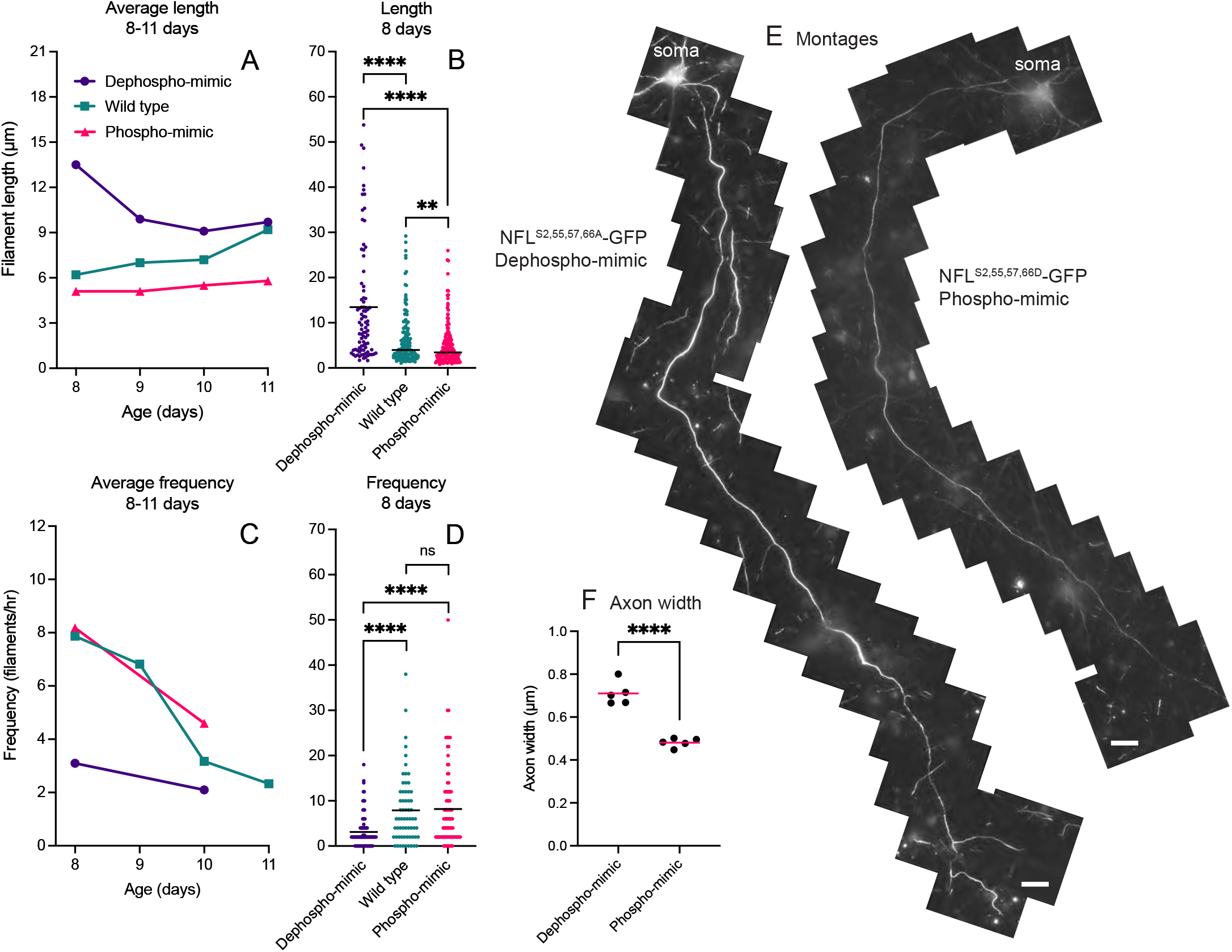
NFL phospho- and dephospho-mimics influence neurofilament length, transport frequency and accumulation. Neonatal rat cortical neurons expressing either wild type NFL-GFP, NFL^S2,55,57,66A^-GFP (dephospho-mimic), or NFL^S2,55,57,66D^-GFP (phospho-mimic). **A.** Average neurofilament length versus age in culture. Each data point represents the average of 87-253 filaments (average=163) measured in 4-9 time-lapse movies (average=5.9) of 15 min duration. **B.** Dot plot of neurofilament lengths at 8 days. We measured 87 filaments in 5 movies for the dephosphomimic, 143 filaments in 5 movies for the wild type, and 146 filaments in 5 movies for the phospho-mimic. **C.** Average neurofilament transport frequency versus age in culture. Each data point represents the average of 28-360 filaments (average=144) measured in 21-88 axons (average=52) in 3-14 time-lapse movies (average=8) of 30 min duration. **D.** Dot plot of neurofilament transport frequencies at 8 days. We measured 131 filaments in 86 axons in 10 movies for the dephospho-mimic, 248 filaments in 63 axons in 13 movies for the wild type, and 360 filaments in 88 axons in 11 movies for the phospho-mimic. The horizontal black lines represent the population means. Note that in A-D there is an age-dependent increase in filament length accompanied by an age-dependent decrease in transport frequency in neurons expressing wild type NFL-GFP but at all ages the filaments were longer and moved less frequently in neurons expressing the dephospho-mimic and were shorter in neurons expressing the phospho-mimic. For the dephospho-mimic, filament length was significantly lower on day 10 compared to day 8 (p=0.0042) but was not significantly different between day 8 and 9 (p=0.10), or between day 8 and 11 (p=0.08). In the phospho-mimic, filament length on day 10 (p=0.041) and 11 (p=0.023) were significantly greater than on day 8. For the wild type, filament length was significantly greater on day 11 compared to days 8 (p<0.0001), 9 (p=0.05), and 10 (p=0.0034). Filaments containing the dephospho-mimic were significantly longer than filaments containing the phospho-mimic on days 8 (p<0.0001), 9 (p<0.0001), 10 (p<0.0001), and 11 (p<0.0001). Filaments containing the dephospho-mimic were significantly longer than filaments containing the wild type on days 8 (p<0.0001), 9 (p=0.0047), 10 (p=0.0034), but not on day 11 (p=0.1). Filaments containing the phospho-mimic were significantly shorter than for the wild type on days 8 (p=0.01), 9 (p<0.0001), 10 (p=0.0061), and 11 (p<0.0001). **E.** Montages of axons of cortical neurons transfected with the dephospho-mimic NFL^S2,55,57,66A^-GFP or the phospho-mimic NFL^S2,55,57,66D^-GFP and then imaged live 16-21 days later using the spinning disk confocal microscope in widefield bypass mode. Scale bars=25 μm. **F.** Average axon width for axons expressing NFL^S2,55,57,66A^-GFP dephospho-mimic (n=5) or NFL^S2,55,57,66D^-GFP phospho-mimic (n=5). The data were compared using an unpaired t-test for equal variances (p<0.0001). The horizontal magenta lines represent the population means. Note the axon expressing the dephospho-mimic is brighter and wider, indicating a higher neurofilament content.

To investigate the mechanism by which NFL phosphorylation influences neurofilament length, we acquired multi-field time-lapse movies and quantified the frequency of neurofilament severing and annealing (Fig. 6). Filaments containing the dephospho-mimic or phospho-mimic constructs exhibited length-dependent severing and annealing as described above (Fig. 6A-D) and the magnitude of this dependence was not significantly different between the dephospho- and phospho-mimics (p=0.16 for severing and p=0.23 for annealing). For every μm increase in length, the probability of severing increased by 3.4% (95% confidence interval: 1.01-1.06, p=0.0063) and the probability of annealing decreased by 4.5% (95% confidence interval: 0.92-0.99, p=0.012). The probability of annealing was also not significantly different between the dephospho-mimic (average frequency=0.74 events/filament/hour, n=122 filaments) and the phospho-mimic (average frequency=0.76 events/filament/hour, n=122 filaments) (p=0.41). However, the probability of severing was 90% less for the dephospho-mimic (average frequency=0.16 events/filament/hour, n=122 filaments) compared to the phospho-mimic (average frequency=1.23 events/filament/hour, n=122 filaments) (95% confidence interval: 0.03-0.28, p<0.0001) (Fig. 6E). The lack of a significant difference in the annealing frequencies for the dephospho- and phospho-mimics may reflect the fact that annealing requires an encounter between two filament ends, which is dependent on the frequency of movement. Thus, it is possible that any decrease in the tendency of filaments containing the phospho-mimic to anneal could be masked by an increased in their frequency of movement, which would result in an increase in end-to-end encounters. Nevertheless, our data are consistent with the hypothesis that NFL phosphorylation favors neurofilament severing.

**Fig. 6.**
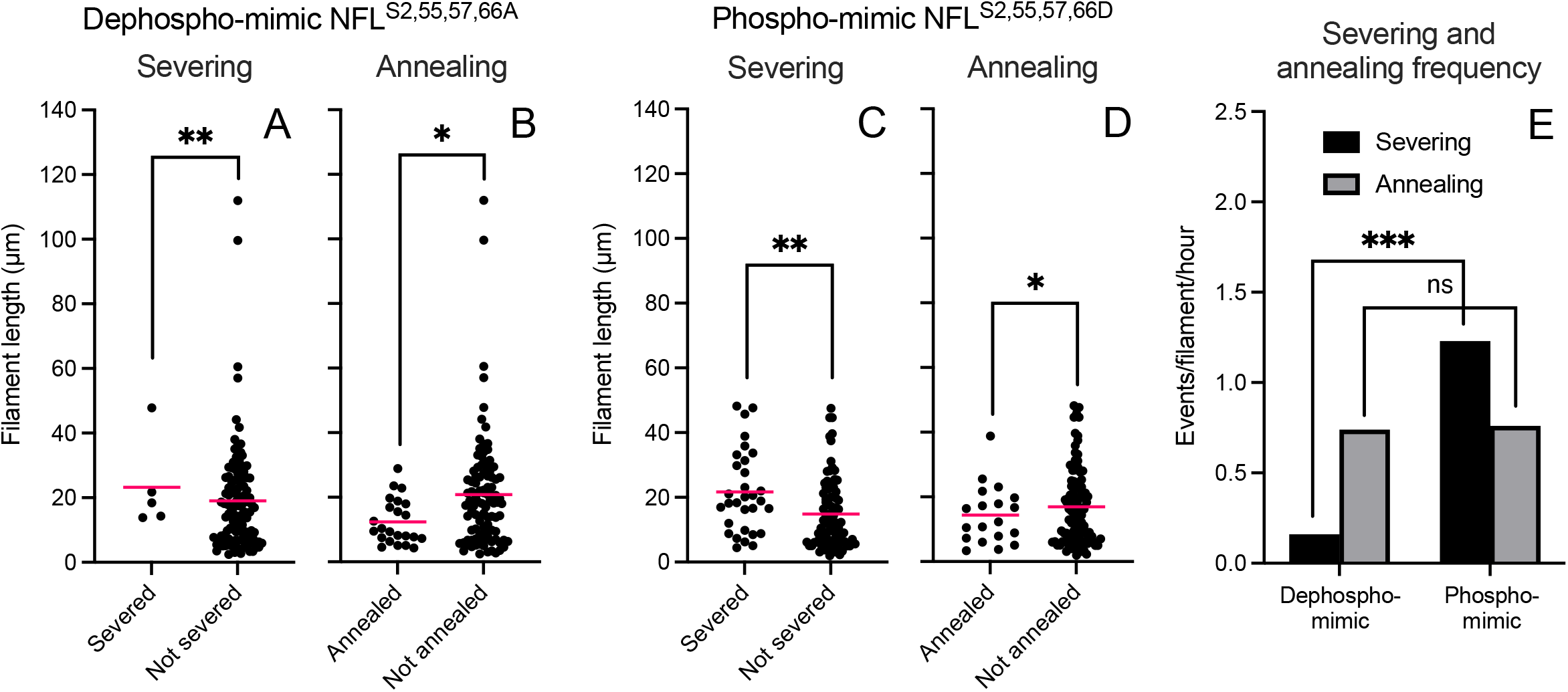
NFL phospho- and dephospho-mimics alter neurofilament annealing and severing frequency. Data from time-lapse movies of neonatal rat cortical neurons expressing either NFL^S2,55,57,66A^-GFP (dephospho-mimic) or NFL^S2,55,57,66D^-GFP (phospho-mimic) after 8-10 days in culture. **A-D.** Dot plots of filament length for filaments that either annealed or did not anneal and that either severed or did not sever during the period of observation. Data from 122 filaments in 73 movies for the dephospho-mimic (total observation time=31.2 hr; average time-lapse movie duration=25.6 min) and 122 filaments in 68 movies for the phospho-mimic (total observation time=25.2 hr; average time-lapse movie duration=22.5 min). Note that there was no significant difference between the length dependence of severing and annealing in the presence of the phospho- and dephospho-mimics. For both mutants, long filaments severed more frequently than short filaments (p=0.0063 in A and C) and short filaments annealed more frequently than long filaments (p=0.012 in B and D). The horizontal magenta lines represent the population means. Note that in the presence of the dephospho-mimic, we encountered four filaments that were longer than 50 μm (specifically 57, 61, 100, and 112 μm). **E.** The average severing and annealing frequencies for the data in A-D. Note that severing is enhanced in filaments containing the phospho-mimic compared to the dephospho-mimic (p<0.0001), whereas annealing frequency is unaffected (p=0.407).

### Activators of protein kinase A reduce neurofilament length

Three of the four sites mutated in the phospho-mimic and dephospho-mimic NFL constructs used above are substrates for protein kinase A (PKA) (Cleverley et al., 1998; Giasson et al., 1996; Giasson and Mushynski, 1998; Hisanaga et al., 1994; Nakamura et al., 2000; Sihag et al., 2007; Sihag and Nixon, 1990; Sihag and Nixon, 1991). Phosphorylation by PKA *in vitro* causes local thinning and fragmentation of NFL homopolymers and native neurofilaments at sites along their length (Hisanaga et al., 1994), and also arrests the assembly NFL homopolymers at the octamer stage (Hisanaga et al., 1990). The neurofilament head domains can also be dephosphorylated by protein phosphatases 1 and 2A (PP1 and PP2A, respectively) (Sacher et al., 1994; Strack et al., 1997). Protein phosphatase 2A (PP2A) dephosphorylates neurofilament proteins that have been phosphorylated by PKA and renders them competent to assemble (Giasson et al., 1996; Saito et al., 1995). Okadaic acid, which is a selective inhibitor of PP1 and PP2A, increases neurofilament head domain phosphorylation and induces reversible neurofilament fragmentation and disassembly in cultured neurons (Giasson et al., 1996; Giasson and Mushynski, 1998; Sacher et al., 1992; Sacher et al., 1994). This effect is thought to be due primarily to PP2A inhibition (Sacher et al., 1994) and is potentiated by PKA activation (Giasson et al., 1996). Collectively, these findings demonstrate that the structural integrity of neurofilaments is regulated by phosphorylation and dephosphorylation of the neurofilament protein head domains, particularly by the antagonistic effects of PKA and PP2A. Thus, we hypothesized that phosphorylation by PKA is a potential mechanism for neurofilament severing in axons.

To test whether PKA regulates neurofilament length in neurons, we treated cultured neonatal rat cortical neurons expressing wild type or dephospho-mimic NFL^S2,55,57,66A^-GFP with 2 mM 8-bromoadenosine 3’,5’-cyclic monophosphate (8-Br-cAMP), which is a membrane-permeant activator of PKA, and/or 5 nM okadaic acid after 9 days in culture. We then measured the lengths of moving neurofilaments in time-lapse movies as described above (Fig. 7). The average filament length was 10.7±9.2 μm in untreated neurons expressing NFL-GFP (Fig. 7A). 8-Br-cAMP reduced the average length to 9.1±7.6 μm (p=0.0085). The average filament length in neurons treated with okadaic acid was 8.1±6.5 μm, but this was not significantly different from the untreated control (p=0.057). However, a combination of okadaic acid plus 8-Br-cAMP reduced the average length to 5.0±3.7 μm (p<0.0001). To test if this effect was due to a direct action on neurofilaments, we repeated these experiments with neurons expressing NFL^S2,55,57,66A^-GFP, which is not phosphorylatable at these four sites (Fig. 7B). The average filament length was 12.9±11.2 μm. As expected, this was significantly greater than in neurons expressing NFL-GFP (p=0.0001). 8-Br-cAMP reduced the average length to 10.4±8.3 μm (p=0.037). The average filament length in neurons treated with okadaic acid was 12.6±8.1 μm, which was not significantly different from the untreated control (p=0.56). A combination of okadaic acid plus 8-Br-cAMP reduced the average length to 8.2±7.6 μm (p<0.0001), but the magnitude of this reduction was less than for GFP-NFL (p<0.0001). Thus, transfection of the neurons with dephospho-mimic NFL^S2,55,57,66A^-GFP partially protected the neurofilaments from the effects of PKA activation.

**Fig. 7.**
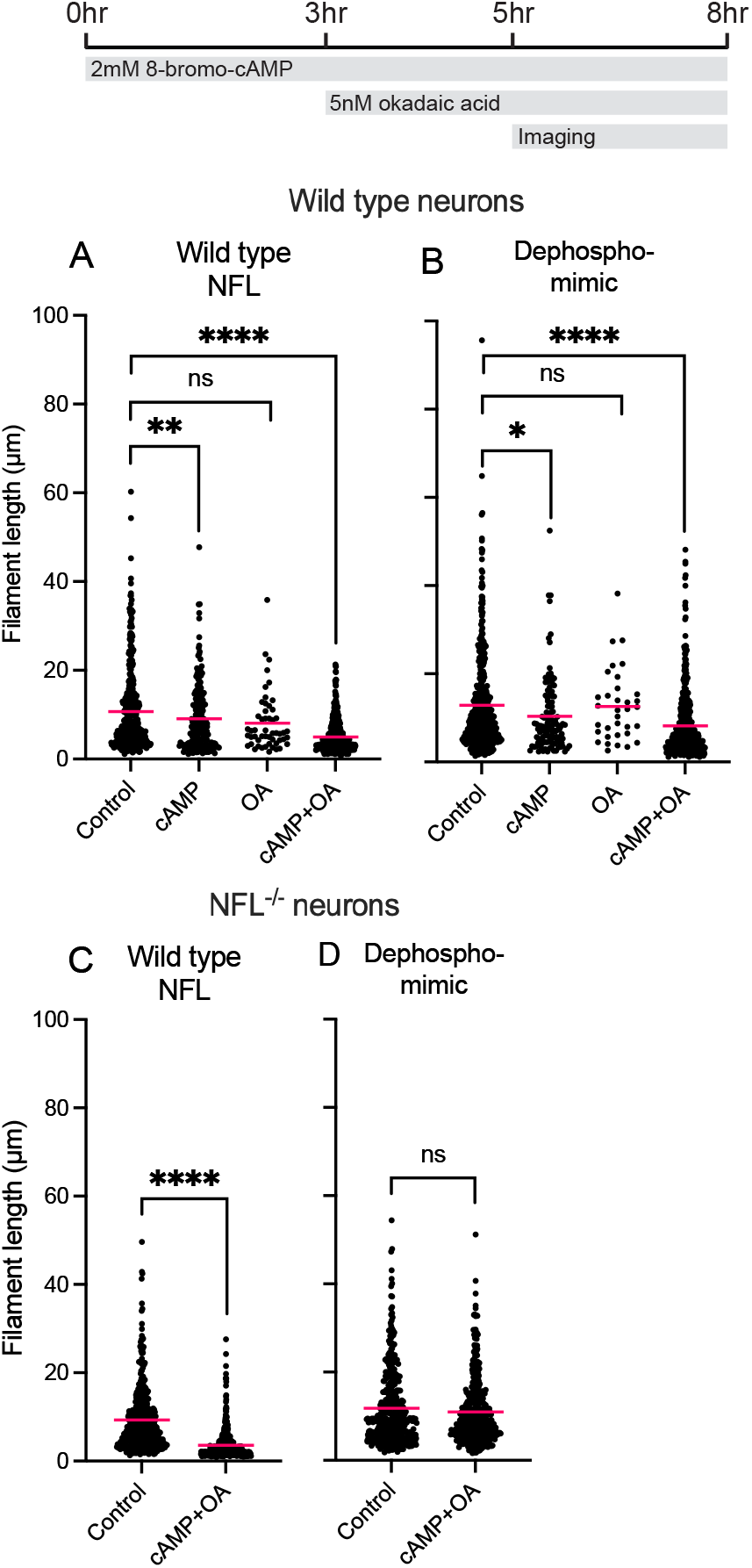
NFL dephospho-mimic protects neurofilaments against the effect of protein kinase A activators. **A,B.** Dot plots of neurofilament length in neonatal rat cortical neurons after 12-14 days in culture (148 movies, average duration=15 min, total of 2,212 neurofilaments tracked). The neurons were transfected with wild type rat NFL-GFP (A) or the dephospho-mimic rat NFL^S2,55,57,66A^-GFP (B) and treated with vehicle, 8-bromo-cAMP (cAMP), okadaic acid (OA), or 8-bromo-cAMP plus okadaic acid. Note that treatment with cAMP but not okadaic acid reduced neurofilament length in neurons expressing wild type NFL, and the effect was more marked for neurons treated with cAMP and okadaic acid together. The same treatments also reduced neurofilament length in neurons expressing the dephospho-mimic, but the extent of the reduction was less. **C,D.** Dot plots of neurofilament length in neonatal mouse cortical neurons from NFL knockout mice after 8-10 days in culture (77 movies, average duration=15 min, total of 1,768 neurofilaments tracked). The neurons were transfected with wild type NFL-GFP (C) or the dephospho-mimic NFL^S2,55,57,66A^-GFP (D) and then treated with vehicle or 8-bromo-cAMP and okadaic acid. The schematic above the plots shows the experimental timeline, which involved pre-treatment for 3 hr with 8-bromo-cAMP, treatment for 2 hr in the presence of 8-bromo-cAMP plus okadaic acid, and imaging for up to 3 hr in the continued presence of these reagents. Note that the treatment reduced neurofilament length in neurons expressing wild type NFL, but not in neurons expressing the dephospho-mimic.

A possible explanation for why the dephospho-mimic NFL^S2,55,57,66A^-GFP offered only partial protection against activators of PKA is that the cells expressed endogenous phosphorylatable NFL in addition to the exogenous mutant NFL. To address this, we repeated the experiment in cortical neurons from neonatal NFL^-/-^ knockout mice using mouse constructs of NFL-GFP and NFL^S2,55,57,66A^-GFP. As expected, the average filament length was greater (11.8±8.6 μm) in neurons expressing the dephospho-mimic NFL^S2,55,57,66A^-GFP than in neurons expressing NFL-GFP (9.3±7.2 μm) and this difference was statistically significant (p<0.0001). In addition, the protective effect of this mutant was much more pronounced. Treatment of cortical neurons expressing NFL-GFP with 8-Br-cAMP plus okadaic acid reduced the average filament length from 9.3±7.2 μm to 3.6±3.4 μm (p=<0.0001) (Fig. 7C). In contrast, the same treatment of cortical neurons expressing NFL^S2,55,57,66A^-GFP, which is not phosphorylatable at these four sites, resulted in only a slight reduction in neurofilament length from 11.8±8.6 μm to 11.0±7.3 μm, which was not significant (p=0.49) (Fig. 7D). Thus, the dephospho-mimic NFL^S2,55,57,66A^-GFP increased filament length and protected the neurofilaments from the effects of PKA activation. These data indicate that the effect of PKA activators on neurofilament length was dependent on phosphorylation of NFL and that the kinase acts on one or more of the four sites mutated in the NFL^S2,55,57,66A^-GFP construct.

## DISCUSSION

We have used live-cell imaging to investigate the dynamics of neurofilament severing and end-to-end annealing in cultured neurons. Our data establish that severing and annealing are robust and quantifiable features of neurofilament dynamics in neurons that regulate neurofilament length, which in turn has a profound influence on the long-term transport behavior of these cytoskeletal polymers. Severing creates shorter filaments which move more persistently, pausing and reversing less often. On the other hand, annealing creates longer filaments which pause and reverse direction more often, resulting in less net movement. Our data also reveal a novel dependency of filament severing and annealing on neurofilament length. Long filaments sever more frequently and anneal less frequently, whereas short filaments sever less frequently and anneal more frequently. Thus, neurofilaments undergo a dynamic cycle of severing and end-to-end annealing in axons that regulates neurofilament length and transport (Fig. 8A).

**Fig. 8.**
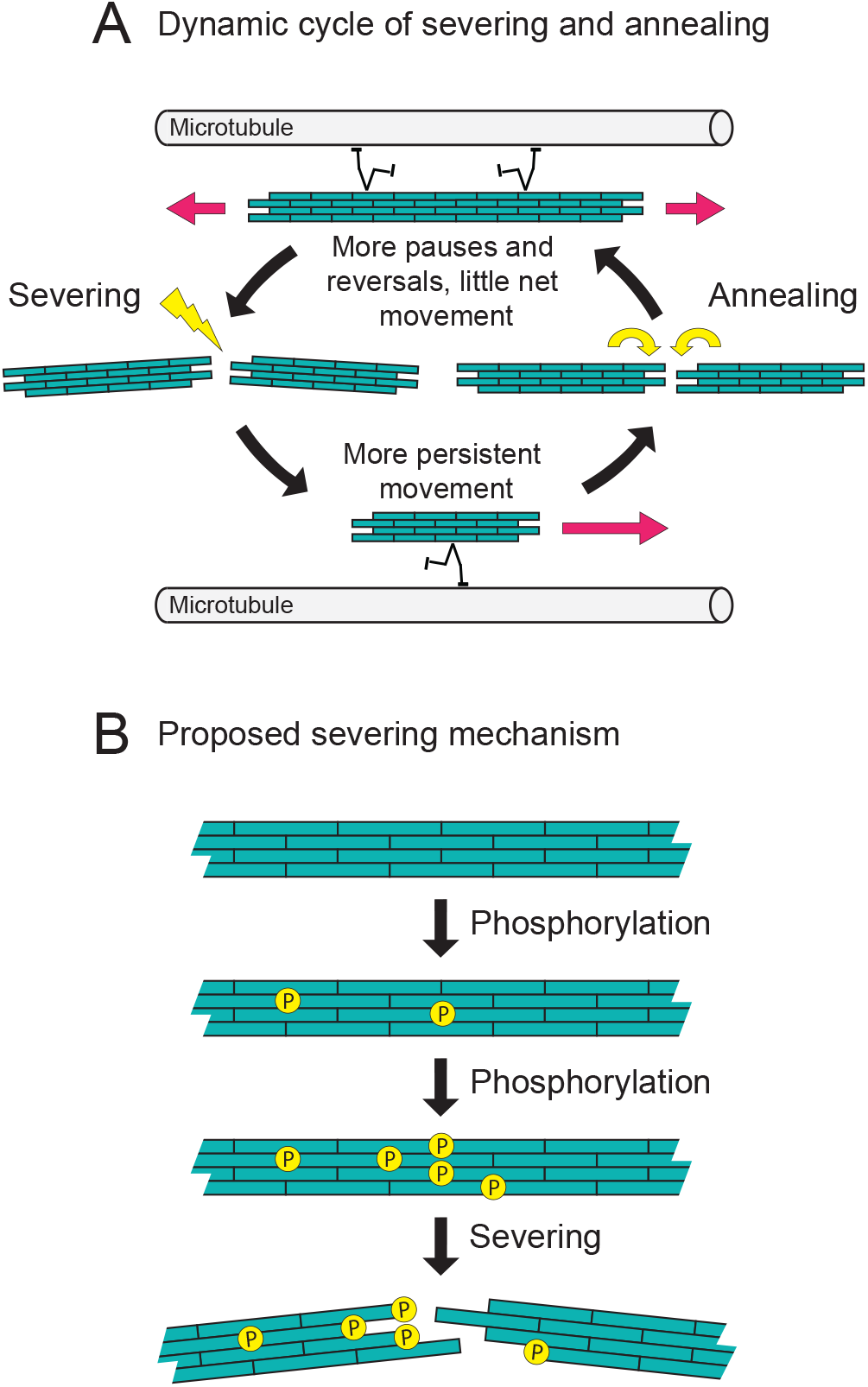
Regulation of neurofilament transport by a dynamic cycle of neurofilament severing and annealing. **A.** Schematic diagram of the cycle of neurofilament severing and annealing described in this study. Neurofilaments move along microtubule tracks (magenta arrows). Severing generates shorter filaments that move more persistently in either the anterograde or retrograde direction, pausing and reversing less often. Annealing generates longer filaments, which pause and reverse direction more often, resulting in little net movement. Short filaments anneal more frequently than long filaments, whereas long filaments sever more frequently than short filaments. The result is a dynamic cycle of severing and annealing that regulates neurofilament transport. **B.** Proposed severing mechanism. Phosphorylation of subunits in the neurofilament polymer weakens their affinity for each other. When the number of phosphorylated subunits reaches a certain critical load, it can destabilize the polymer at that location and cause the polymer to break.

According to this model, neurofilament length is determined by the balance of the annealing and severing rates. At steady state, which may be achieved in mature non-growing axons, the two rates would be balanced. In our cultures, where the average neurofilament length increased with time, we thus expected to observe more annealing than severing in our time-lapse movies. However, we observed about twice as many severing events as annealing events. A possible explanation for this is that we can only detect annealing and severing in axons that have very low neurofilament content, where single filaments can be observed in isolation, yet annealing requires an encounter between two filament ends and thus may be less likely when neurofilament density is low. In contrast, severing does not require a diffusional encounter between filaments and thus may be independent of filament density. Thus, our observations in these cultured neurons may underestimate of the rate of annealing that would be encountered in more neurofilament-rich axons.

The mechanism of the length-dependence of neurofilament movement, severing and annealing is an interesting question. The length-dependence of the pauses and reversals could be explained if the number of motors recruited by the filaments is proportional to the filament length. Short filaments might recruit on average one motor, leading to persistent unidirectional movement, whereas long filaments might recruit on average multiple motors that could be of opposite directionality. The latter would result in a sort of dynamic tug-of-war that might cause long filaments to stall for prolonged periods or shuttle between alternating anterograde and retrograde bouts, resulting in little net movement. With respect to the length-dependence of neurofilament severing and annealing. it is possible that long filaments anneal less frequently simply because they are less mobile and thus collide less frequently and that they sever more frequently simply because longer polymers have more potential severing sites. However, it is also possible that these differences between long and short filaments reflect differences in their composition, post-translational modification or interactions with other cytoplasmic components.

To investigate the effect of phosphorylation on neurofilament length and transport, we used site-directed mutagenesis. Expression of the phospho-mimic mutant (NFL^S2,55,57,66D^) resulted in shorter neurofilaments that moved and severed more frequently, whereas the non-phosphorylatable mutant (NFL^S2,55,57,66A^) had the opposite effect. Remarkably, filaments containing the non-phosphorylatable mutant reached lengths of up to 96 μm, which is the longest moving filament we have ever encountered by live imaging of neurons expressing neurofilament proteins. Treatment with activators of protein kinase A reduced neurofilament length and this effect was blocked by mutating serines 2, 55, 62 and 66 in the NFL head domain to alanines. These observations suggest that N-terminal phosphorylation of NFL could be a mechanism for neurofilament severing. Consistent with this severing model, neurofilaments containing the dephospho-mimic exhibited lower rates of severing, while filaments containing the phospho-mimic exhibited higher rates of severing.

Since neurofilaments contain about 32 polypeptides per cross-section (Heins et al., 1993), severing likely requires the phosphorylation of multiple subunit proteins at the same location along the polymer, with each additional phosphorylated subunit further weakening the filament at that location and increasing the severing probability (Fig. 8B). While the phosphorylation could be random, it is more likely to occur locally downstream of local signaling events that may be spatially or temporally controlled to regulate neurofilament length and transport. Severing cannot be generated simply by the pulling force of molecular motors because it takes nanonewtons of force to stretch and break single intermediate filaments (Guzman et al., 2006; Kreplak et al., 2005; Kreplak et al., 2008), but it is possible that such a pulling force could break a filament that is weakened by local phosphorylation events. In this regard, it is interesting to note that the severing events in our movies often appear to occur when the filaments begin to move. However, since severing cannot be confirmed until the daughter fragments move apart, we cannot rule out the possibility that these filaments severed spontaneously prior to the onset of movement.

We focused on PKA in this study because it has been reported to phosphorylate three of the sites mutated in our phospho- and dephospho-mimic NFL constructs. However, we do not exclude the possible involvement of other kinases and/or neurofilament subunits. PKA can phosphorylate the head domains of peripherin, NFM and NFH in addition to NFL (Cleverley et al., 1998; Giasson and Mushynski, 1998; Hisanaga et al., 1994; Huc et al., 1989; Sihag et al., 1999), and the NFL head domains can be phosphorylated by protein kinase C (PKC), Rho kinase, protein kinase N (PKN), and CaMKII (Gonda et al., 1990; Hashimoto et al., 1998; Hashimoto et al., 2000; Hisanaga et al., 1994; Mukai et al., 1996; Nakamura et al., 2000; Sihag et al., 2007; Sihag and Nixon, 1991). PKC has also been reported to phosphorylate NFM and NFH (Sihag et al., 1988) and has been shown to induce intermediate filament disassembly *in vitro* and in cultured cells (Giasson et al., 1996; Hisanaga et al., 1990; Hisanaga et al., 1994). Nonetheless, our data indicate that treatment with cAMP and okadaic acid is sufficient to induce neurofilament severing, and this can be rescued by expression of the non-phosphorylatable NFL mutant in NFL knockout neurons. Thus, PKA phosphorylation appears to be sufficient to induce severing, and NFL appears to be the principal target of this kinase. Since head domain phosphorylation by PKA also induces disassembly of vimentin and keratin filaments (Ando et al., 1989; Ando et al., 1996; Eriksson et al., 2004; Evans, 1988; Inagaki et al., 1987), focal destabilization of intermediate filaments by N-terminal phosphorylation of their constituent polypeptides at specific locations along their length may be a general mechanism for severing this class of cytoskeletal polymers. In this way, phosphorylation at discrete sites along intermediate filaments may represent another tool in the toolbox that cells can use to regulate the organization and dynamics of this class of cytoskeletal polymers.

The axonal transport of neurofilaments is known to slow both spatially along axons and temporally during post-natal development of myelinated axons (Hoffman et al., 1985; Hoffman et al., 1983; Jung and Brown, 2009; Xu and Tung, 2001). This contributes to the accumulation of neurofilaments that drives the expansion of myelinated axon caliber (Cleveland et al., 1991; Hoffman, 1995). The mechanism of this slowing has been attributed to an increase in phosphorylation of the carboxy-terminal tail domains of neurofilament proteins M and H (Ackerley et al., 2003; Garcia et al., 2003; Rao et al., 2003; Sanchez et al., 2000). However, deleting these domains or mutating them to block phosphorylation did not impair neurofilament transport or accumulation *in vivo* (Garcia et al., 2009; Rao et al., 2002; Yuan et al., 2006). Thus, the role of neurofilament tail domain phosphorylation in the regulation of neurofilament transport is presently unclear. Based on the data presented above, we speculate that an additional or alternative mechanism for slowing neurofilament transport may be the modulation of neurofilament length by spatial and temporal regulation of the cycle of neurofilament severing and annealing (e.g. by modulation of the activities of PKA and PP2A). For example, a shift in these activities to favor annealing over severing during axonal maturation would result in an increase in average neurofilament length and a decrease in net transport velocity. In this way, neurofilament severing and annealing may represent a mechanism to regulate the neurofilament content and diameter of axons, which is an important influence on their conduction velocity.

## ACKNOWLEDGMENTS

We thank Chris Miller (King’s College, University of London) for sharing the rat phospho- and dephospho-mimic NFL constructs, Robert Evans (University of Colorado Denver) for providing the SW13vim-cell line, Don Cleveland (University of California San Diego) for providing the NFL knockout mice originally generated by Jean-Pierre Julien (Université Laval), and Sontoria King for comments on the manuscript. This work was supported by National Institutes of Health Grants R01 NS038526, P30 NS104177, and S10 OD010383 to AB, and by a grant from the Takeda Science Foundation to AU. The authors declare that they have no competing financial interests.

## MOVIE LEGENDS

**Movie S1. Long-range multi-field tracking of a neurofilament**

Multi-field time-lapse movie of a cortical neuron expressing GFP-NFM, showing a short neurofilament (length=4.3 μm) that moves retrogradely along the axon, eventually entering the cell body. The filament is to the lower left of the center of the field of view at the start of the movie. Note that the filament moves persistently in one direction, pausing infrequently, and that it is not visible at 01:48 to 2:03 and 04:14 to 04:45 because the axon passes out of the plane of focus as it courses over other cells at these locations. At 02:03, the filament passes another short filament that is pausing. At 03:06, the filament passes a short filament that is moving in the opposite (anterograde) direction. At 05:09, the filament passes a pausing filament, which subsequently moves in the opposite (anterograde) direction. The movie ends at 06:09 when the filament enters the cell body. Dimensions of field: 318 x 207 μm. Proximal is right, distal is left. Time-lapse interval: 3 sec. Time stamp min:sec. Time compression 21:1.

**Movie S2. An annealing event**

Excerpt from a multi-field time-lapse movie of a cortical neuron expressing GFP-NFM, showing a long neurofilament (length=20.8 μm) that moves anterogradely initially. The filament is in the center of the field of view at the start of the movie. The filament pauses at 03:00 and then anneals at its proximal end with a short retrogradely moving neurofilament (length=6.3 μm). The resulting daughter filament (length=27.5 μm) then shuttles back and forth repeatedly from 03:00 to 09:00, as if engaged in a tug-of-war, before moving persistently in a retrograde direction. Dimensions of field: 171 x 170 μm. Proximal is upper left, distal is lower right. Time-lapse interval: 3 sec. Time stamp min:sec. Time compression 21:1.

**Movie S3. A severing event**

Excerpt from a multi-field time-lapse movie of a cortical neuron expressing GFP-NFM, showing a long neurofilament (length=29 μm). The filament is in the center of the field of view at the start of the movie. The filament moves anterogradely in a folded configuration initially, fully unfolding at 03:51. The filament then appears to sever at 04:15, though this is not clearly evident until the two daughter filaments separate at 05:21. The shorter (proximal) daughter filament (length=7.1 μm) moves away in a retrograde direction, then the longer (distal) daughter filament (length=21.6 μm) folds up on itself at 05:36 before unfolding and resuming anterograde movement at 07:12. Dimensions of field: 118 x 133 μm. Proximal is lower left, distal is upper right. Time-lapse interval: 3 sec. Time stamp min:sec. Time compression 21:1.

**Movie S4. Rapid persistent movement of a short neurofilament**

Multi-field time-lapse movie of a cortical neuron expressing GFP-NFM, showing a short neurofilament (length=5.4 μm) that moves anterogradely. The filament is to the right of the center of the field of view at the start of the movie. It briefly folds and unfolds at 02:03 to 02:09, and then at 02:36 it catches up with a shorter filament (length=2.3 μm) that is also moving anterogradely and the two filaments anneal. The daughter filament (length=7.6 μm) continues to move anterogradely, passing another pausing filament at 03:18 to 03:30 and then at 04:24 it pauses. At 04:30 a long filament moving retrogradely overlaps the short filament and then pauses, at which point the short filament can no longer be resolved. Note that the movie drifts out of focus from 03:51 to 04:27. Dimensions of field: 347 x 101 μm. Proximal is left, distal is right. Time-lapse interval: 3 sec. Time stamp min:sec. Time compression 21:1.

**Movie S5. Frequent pausing and reversal of a long neurofilament**

Multi-field time-lapse movie of a cortical neuron expressing GFP-NFM, showing a long neurofilament (length=27.9 μm) that severs twice. The filament is in the center of the field of view at the start of the movie, partially overlapping one or more pausing neurofilaments. The filament moves past these pausing filaments in a net retrograde direction, frequently pausing and reversing direction. Note that the filament remains fully extended during these bidirectional movements, as if pulled alternately at one end and then the other (Fenn et al., 2018b), and that it passes several shorter filaments moving in the opposite (anterograde) direction around 04:42 and 08:51. The filaments occasionally pass out of the plane of focus at locations where the axon crosses another cell. At around 10:06, the filament begins to move persistently back in the anterograde direction, passing several other pausing filaments starting at 13:57 and eventually pauses at 15:30 close to where it started and partially overlapping another pausing filament. At 17:39 the latter filament appears to anneal with a very short filament moving anterogradely and then the daughter filament moves away in the anterograde direction. At 18:30, the filament of interest folds back on itself. Two shorter filaments moving in the retrograde direction then overlap the filament of interest at 20:09 and 21:03, pausing and then moving away at 21:06 and 21:33, respectively. At some point during this period of folding and overlap, the filament of interest severs twice. This becomes evident when the daughter fragments move apart in the anterograde direction at 21:42 and 22:24, giving rise first to three fragments (lengths=10.6, 6.0 and 11.0 μm) that can be resolved from each other in the final frame. Dimensions of field: 310 x 166 μm. Proximal is right, distal is left. Time-lapse interval: 3 sec. Time stamp min:sec. Time compression 21:1.

**Movie S6. Movement of neurofilaments expressing the NFL dephospho-mimic**

Fixed-field time-lapse movie of cortical neurons expressing the NFL^S2,55,57,66A^-GFP dephospho-mimic. Note that the fusion protein co-assembles efficiently into neurofilaments with no detectable diffuse (unassembled) protein. On average, the filaments are longer and move less frequently than in neurons expressing wild type NFL-GFP (Movie S7). See Fig. 5 for quantification. Dimensions of field: 82 x 82 μm. Time-lapse interval: 3 sec. Time stamp min:sec. Time compression 21:1.

**Movie S7. Movement of neurofilaments expressing wild type NFL**

Fixed-field time-lapse movie of cortical neurons expressing wild type NFL-GFP. On average, the filaments are shorter and move more frequently than in neurons expressing the NFL^S2,55,57,66A^-GFP dephospho-mimic (Movie S6). On average, the filaments are longer and move less frequently than in neurons expressing the NFL^S2,55,57,66D^-GFP phospho-mimic (Movie S8). See Fig. 5 for quantification. Dimensions of field: 82 x 82 μm. Time-lapse interval: 3 sec. Time stamp min:sec. Time compression 21:1.

**Movie S8. Movement of neurofilaments expressing the NFL phospho-mimic**

Fixed-field time-lapse movie of cortical neurons expressing the NFL^S2,55,57,66D^-GFP phosphomimic. Note that the fusion protein co-assembles efficiently into neurofilaments with no detectable diffuse (unassembled) protein and no aggregation. On average, the filaments are shorter and move more frequently than in neurons expressing wild type NFL-GFP (Movie S7). See Fig. 5 for quantification. Dimensions of field: 82 x 82 μm. Time-lapse interval: 3 sec. Time stamp min:sec. Time compression 21:1.

**Movie S9. Movement of neurofilaments expressing wild type NFL treated with cAMP and okadaic acid**

Fixed-field time-lapse movie of NFL^-/-^ cortical neurons expressing wild type NFL-GFP and treated with 8-bromo-cAMP and okadaic acid. On average, the filaments are shorter and move more frequently than in neurons expressing the non-phosphorylatable NFL^S2,55,57,66A^-GFP dephospho-mimic and subjected to the same treatment (Movie S10). See Fig. 7 for quantification. Dimensions of field: 82 x 82 μm. Time-lapse interval: 3 sec. Time stamp min:sec. Time compression 21:1.

**Movie S10. Movement of neurofilaments expressing the NFL dephospho-mimic treated with cAMP and okadaic acid**

Fixed-field time-lapse movie of NFL^-/-^ cortical neurons expressing the non-phosphorylatable NFL^S2,55,57,66A^-GFP dephospho-mimic and treated with 8-bromo-cAMP and okadaic acid. On average, the filaments are longer and move less frequently than in neurons expressing wild type NFL-GFP and subjected to the same treatment (Movie S9). See Fig. 7 for quantification. Dimensions of field: 82 x 82 μm. Time-lapse interval: 3 sec. Time stamp min:sec. Time compression 21:1.

